# Functional unknomics: closing the knowledge gap to accelerate biomedical research

**DOI:** 10.1101/2022.06.28.497983

**Authors:** Joao Rocha, Satish Arcot Jayaram, Tim J. Stevens, Nadine Muschalik, Rajen D. Shah, Sahar Emran, Cristina Robles, Matthew Freeman, Sean Munro

## Abstract

The human genome encodes ∼20,000 proteins, many still uncharacterised. Scientific and social factors have resulted in a focus on well-studied proteins, leading to a concern that poorly understood genes are unjustifiably neglected. To address this, we have developed an “Unknome database” that ranks proteins based on how little is known about them. We applied RNAi in *Drosophila* to 260 unknown genes that are conserved between flies and humans. About a quarter are required for viability, and functional screening of the rest revealed hits for fertility, development, locomotion, protein quality control and resilience to stress. CRISPR/Cas9 gene disruption validated a component of Notch signalling and two genes contributing to male fertility. Our work demonstrates the importance of poorly understood genes, provides a resource for future research acceleration, and highlights a need for our awareness of ignorance to be protected from erosion by automated database annotation.

## Introduction

The advent of genome sequencing revealed in humans and other species thousands of genes encoding proteins that had not been identified by previous biochemical or genetic studies. Since the release of the first draft of the human genome sequence in 2000, the function of many of these new proteins has been identified. However, despite over twenty years of extensive effort there are also many others that still have no known function. The mystery and the potential biological significance of these unknown genes is enhanced by many of them being well conserved, and many of them being completely unrelated to known proteins and thus lacking clues to their function. Analysis of publication trends has revealed that research efforts continue to focus on genes and proteins of known function, with similar trends seen in gene and protein annotation databases (Edwards et al., 2011; Peña-Castillo and Hughes, 2007; Sinha et al., 2018). This is despite clear evidence from studies of gene expression and genetic variation that many of the poorly characterised proteins are linked to disease, including those that are eminently druggable (Oprea et al., 2018; Stoeger et al., 2018).

This bias in biological research toward the previously studied appears to reflect several linked factors. Clearly, funding and peer-review systems are more likely to support research on proteins with prior evidence for functional or clinical importance; individual perception of project risk, and social aspects of the research enterprise no doubt also contribute. In addition, scientific factors have been proposed, including a lack of specific reagents like antibodies or small molecule inhibitors, and a tendency to focus on proteins which are abundant and widely expressed and so likely to be present in cell lines and model organisms (Edwards et al., 2011; Stoeger et al., 2018). Finally, some genes may have roles that are not relevant to laboratory conditions (Peña-Castillo and Hughes, 2007).

Whatever the reasons, this bias against the unknown is clear and does not appear to be diminishing. This has led to concern that important fundamental or clinical insight, as well as potential for therapeutic intervention, is being missed, and hence the launch of several initiatives to address the problem. These include programs to generate proteome-wide sets of reagents such as antibodies or mouse knock-out lines (Muñoz-Fuentes et al., 2018; Thul et al., 2017). In addition, the NIH’s Illuminating the Druggable Genome initiative supports work on understudied kinases, ion channels and GPCRs (Rodgers et al., 2018). There have been initiatives to develop new means to predict protein function or structure (*9*-*13*). Finally, databases such as Pharos, Harmonizome and neXtProt link human genes to expression and genetic association studies with the aim of highlighting understudied genes relevant to disease and drug discovery (Duek et al., 2018; Nguyen et al., 2017; Rouillard et al., 2016).

In this work, we have investigated directly the potential biological significance of conserved genes of unknown function by developing a systematic approach to their identification and characterisation. We have created an ‘Unknome database’ that assigns to each protein from a particular organism a ‘knownness’ score based on a user-controlled application of the widely-used Genome Ontology (GO) annotations (Ashburner et al., 2000; The Gene Ontology Consortium, 2019). The database allows selection of an ‘unknome’ for a particular model organism that can be tuned to reflect the degree of conservation in other species, for example allowing a focus on those proteins of unknown function that have orthologs in humans or are highly conserved in evolution. We use this database to evaluate the human unknome and find that it is only slowly shrinking over time. To assess the value of the unknome as a foundation for experimental work we selected a set of 358 *Drosophila* proteins of unknown function that are conserved in humans, and used RNA interference to test their contribution to a wide range of biological processes. This revealed proteins important for diverse biological roles, including cilia function and Notch pathway signalling. Overall, our approach has demonstrated that significant and unexplored biology is encoded in the neglected parts of the human proteome.

## Results

### Construction of an Unknome database

Much of the progress in understanding protein function has come from research in model organisms selected for their experimental tractability. Application of this research to the proteins of humans requires being able to identify the orthologs of these proteins in model organisms. Although it is not certain that orthologs in different species have precisely the same function, they generally have similar or related functions, implying that work from model organisms at the very least provides plausible hypotheses to test. Thus, our Unknome database was designed to link a particular protein with what is known about its orthologs in humans and popular model organisms.

A range of methods for identifying orthologs have been developed based on sequence conservation and although none are perfect, several achieve an accuracy in excess of 70%. We initially used the OrthoMCL database as it covered a wide range of organisms (Fischer et al., 2011). However, OrthoMCL was not being updated, and so the current Unknome database is based on the PANTHER database (version 16.0) which covers over 142 organisms, is currently in continuous development, and has a good level of sensitivity and accuracy (Mi et al., 2021).

The heart of the Unknome database has been the development of an approach to assigning a “knownness” score to proteins. This is not trivial and is inevitably a somewhat subjective measure. Definitions of “known” range from a simple statement of activity to an understanding of mechanism at atomic resolution, and even well characterised proteins can reveal unexpected extra roles. Thus, we designed the database so that the criteria for knownness can be user defined, as well as having a default set of criteria. The Gene Ontology (GO) Consortium provides annotations of protein function that are well suited to this application. Firstly, GO annotation is based on a controlled vocabulary and so is consistent between different species, and secondly, it is well structured thus allowing a user to apply their own definition of knownness.

The Unknome database combines PANTHER protein family groups (which we term “clusters”) with the GO annotations for each member of the cluster. This includes annotations from humans and the 11 model organisms selected by the GO Consortium for their Reference Genome Annotation Project. The sequence-similar protein clusters (primary PANTHER families) not only contain orthologs, but also recent paralogs: duplications within individual species. The knownness score for each protein is based upon the number of GO annotations it possesses.

However, it is important to recognise that GO annotations do not all have equal evidential value, and the Unknome database allows users to account for this in generating a knownness score. This allows application of greater weights to annotations that are more likely to be reliable, such as those from a “Traceable Author Statement” rather than those “Inferred from Electronic Annotation” (Figure 1A). In addition, weighting allows selection of annotations most relevant to function. For instance, a protein’s sub-cellular location is often included in its GO annotation, but this may not helpfully restrict the range of possible functions, so the database provides the option of excluding it when calculating a knownness value. The final knownness score of a cluster of proteins is set as the highest score of a protein in the cluster (Figure 1B). The Unknome database is available as a website (unknome.mrc-lmb.cam.ac.uk) that provides all protein clusters that contain at least one protein from humans or any of the 11 model organisms. The clusters can be ranked by knownness, and the user can modify this list so as to include only those proteins that are present in a particular combination of species, such as human plus a preferred model organism (Figure 1C). For each protein family, the interface shows the orthologs in its cluster and how the knownness of the cluster has changed over time (Figure 1D). These design principles maximise the versatility and power of the Unknome database as a tool for researchers from different biomedical fields.

**Figure 1.**
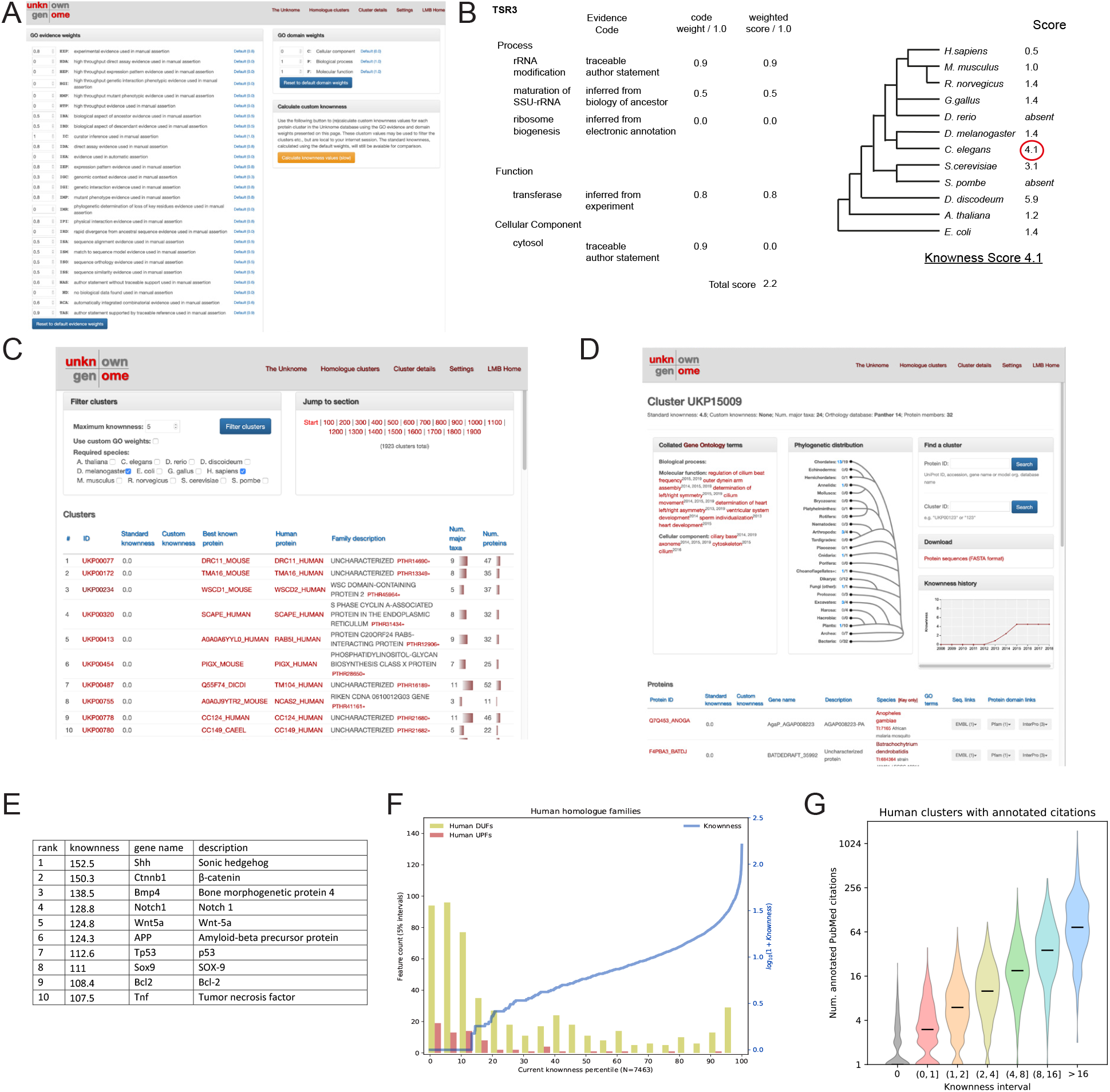
The Unknome database. (A) User interface to weight GO annotations, showing the default weightings we used to generate an unknome gene set. (B) Calculation of a knownness score for a cluster of orthologs based on the highest score in the cluster. (C) User interface to list proteins from select model organisms by the knownness of their ortholog cluster. (D) Information for each cluster showing the distribution across species, links to information for the protein from each species, and the change in knownness over time. (E) The top ten known gene clusters, showing the best known gene in each. All the genes are from mouse with the exception of human APP. (F) Clusters in the unknome that contain at least one human protein ranked by knownness, showing the distribution of proteins that are defined by Pfam as being an unknown protein family (UPF) or containing a domain of unknown function (DUF). (G) Plot of the number of PubMed publications for genes of the indicate range of knownness.

### Validation of the Unknome database

To validate the Unknome database we ranked the 7,463 clusters of orthologs and paralogs that contain at least one human protein, a criterion based on our ultimate goal of revealing new human biology. Validating the overall approach, the top 10 scoring proteins have well known roles in development and cell function (Figure 1E). In contrast, proteins containing one of the “Domains of Unknown Function” defined by the Pfam database were concentrated at the bottom of the range (Figure 1F). Clusters with a score of 1.0 or less correspond to 18.3% of all clusters but to 36% of the DUFs, and 59% of the related Unknown Protein Function (UPFs). The exceptions were typically multidomain proteins of known function which contain one domain whose role is unclear. Finally, the total number of PubMed entries on each protein shows a good correlation with their current knownness (Figure 1G). Overall, we conclude that the calculated knownness score provides a useful means to identify proteins of unknown function.

### The change of the Unknome over time

Unlike most databases, the Unknome will shrink over time. The knownness scores for clusters containing human proteins have increased across the whole range of proteins, but the proportion with a knownness score of 1 or less has only declined from 58% to 39% over the last ten years (Figure 2A). This slow progress reflects the rather narrow research activity on the proteins: human genes and proteins are much more likely to have been published on in the last ten years if they are in clusters that were already well known at the start of this period (Figure 2B). This gives further support to the notion that research activity tends to focus on what has already been studied in depth (Edwards et al., 2011; Pfeiffer and Hoffmann, 2007). There are 726 human genes whose knownness was zero ten years ago but has since increased to above two. The GO terms most enriched in this set are mostly associated with cilia, reflecting recent acceleration of progress in studying this large and complex structure that is absent from some model organisms such as yeast (Figure 2C). Consistent with this, the less known human genes tend to be less likely to be conserved outside of vertebrates, suggesting that progress has been hampered by there being fewer orthologs that could be found by genetic screens in non-vertebrates (Figure 2D). Interestingly, the most highly known proteins are also less likely to be conserved outside of metazoans, reflecting the fact that many are involved in important developmental pathways or signalling events relevant to multicellularity. However, of the 1710 human-containing clusters with a current knownness score of less than 2.0, 47% are detectably conserved outside of vertebrates and 29% are conserved outside of metazoans (Figure 2E). Interestingly, no one model organism contains all of these, indicating that each has a role to play in illuminating the human unknome.

**Figure 2.**
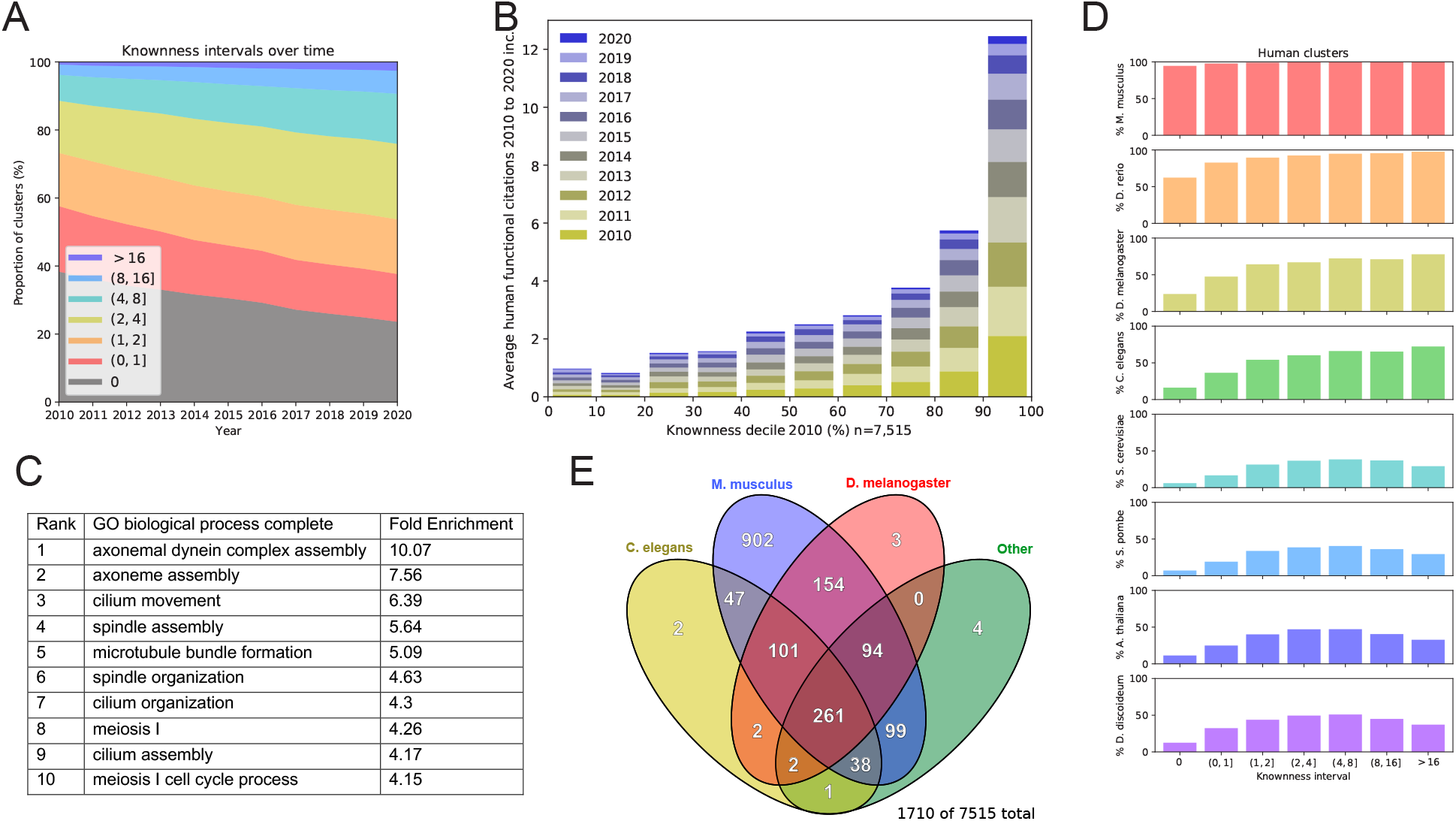
Analysis of trends in knownness. (A) Change in the distribution of knownness of the 7463 clusters that contain at least one human protein. (B) Number of papers to appear in PubMed since 2010 on the 7463 clusters ranked into deciles based on knownness at 2010. The best known papers in 2010 received the most publications in subsequent years. (C) The 10 largest GO term enrichments for the 726 human proteins whose knownness has increased from 0 in 2008 to above 2 in 2018. (D) Conservation in model organisms of human genes clustered by knownness. (E) Venn diagram showing the distribution of genes from the indicated species in the 1842 clusters of knowness <2.0 and which contain at least one human protein.

### Functional unknomics in *Drosophila*

To test the value of the Unknome database, and to pilot experimental approaches to studying neglected but well conserved proteins, we selected a set of unknown human proteins that are conserved in *Drosophila* and hence amenable to genetic analysis. *Drosophila* also lack the partial redundancy between closely related paralogs that arose in many gene families from the two whole genome duplications that occurred early in vertebrate evolution (Holland and Ocampo Daza, 2018). A powerful approach to the function of gene in *Drosophila* is to knock down its expression with RNAi and assess the biological consequences (Homem et al., 2014; Mummery-Widmer et al., 2009). We thus determined the effect of expressing hairpin RNAs to direct RNAi against this panel of genes of unknown function.

We initially selected all genes that had a knownness score of ≤1.0 and are conserved in both humans and flies, as well as being present in at least 80% of available metazoan genome sequences. Of the 629 corresponding *Drosophila* genes, 358 were available in the KK library that was the best available genome-wide RNAi library at the time (Table S1) (Heigwer et al., 2018). These stocks were crossed to lines containing Gal4 drivers to express the hairpin RNAs in either the whole fly or in specific tissues. After testing for viability, the non-essential genes were then screened with a panel of quantitative assays designed to reveal potential roles in a wide range of biological functions. These included male and female fertility, tissue growth (in the wing), response to the stresses of starvation or reactive oxygen species, proteostasis, and locomotion. The results of these screens are discussed below.

### Unknown genes have essential functions

To determine if the genes were required for viability a ubiquitous driver was used to direct RNAi throughout development. For 162 of the 358 genes the resulting progeny showed compromised viability with either all (lethal) or almost all (semi-lethal) failing to pass beyond pupal eclosion, suggesting that these genes are essential for development or cell function (Table S1). However, it was subsequently reported that in a subset of the lines in the KK RNAi library the transgene was in an integration locus (40D) that itself results in serious developmental defects when the transgene is expressed with a GAL4 driver (Green et al., 2014; Vissers et al., 2016). Following PCR screening, we removed all of the stocks that had this integration site, all but one of them having been been lethal in the initial screen. For the remaining 260 genes, the stocks used the alternative integration site which is not problematic, with KK stocks having been used successfully in a range of different screens (Czech et al., 2013; Homem et al., 2014). For these, the RNAi compromised viability in 62 cases (24%). Of these 62 genes, 12% were also identified in a recent genome-wide screen of genes required for viability of S2 cells; in contrast, only 4% of the viable genes were hits in the S2 cell screen (Viswanatha et al., 2018). The S2 study estimated that 17% of genes known to be essential in flies are also essential in S2 cells, and it is likely that using RNAi to knock down gene function underestimates lethality. Our screen in whole organisms reveals that, despite several decades of extensive genetic screens in *Drosophila*, there are many genes with essential roles that have eluded characterisation.

Of course, there is more to life than merely being alive. We therefore subjected the 198 apparently non-essential genes to a range of phenotypic tests to determine if they had detectable roles in a wide range of organismal functions. The results of these function screens are described below, followed by a validation of selected hits.

### Contribution of unknome genes to fertility

To test fertility, specific GAL4 drivers were used to knockdown the set of 198 unknown genes in either the male or female germline. Even with collecting data for multiple flies per gene, the resulting brood sizes showed some variability, as expected for a quantitative measure of a biological process. Thus, for all our assays we needed to determine if outliers had a phenotype that exceeded to a statistically significant degree the variation intrinsic in the population. To do this, we used statistical tests based on three steps. First, we performed a regression on the replicate data for each gene to estimate its parameters and standard errors within the assay. Next, an outlier region was determined by fitting the parameter estimates for all analysed genes to a normal distribution, which was then used to define a boundary for outliers. Finally, for each gene, we tested the hypothesis that it falls within the outlier boundary. This approach is described in detail in the Supplemental Information. To display the data from the fertility tests, mean brood sizes obtained from RNAi-treated males was plotted against those obtained from RNAi-treated females for each gene (Figure 3A). Several of the RNAi lines gave a substantial reduction in brood size that was sex-specific and highly statistically significant.

**Figure 3.**
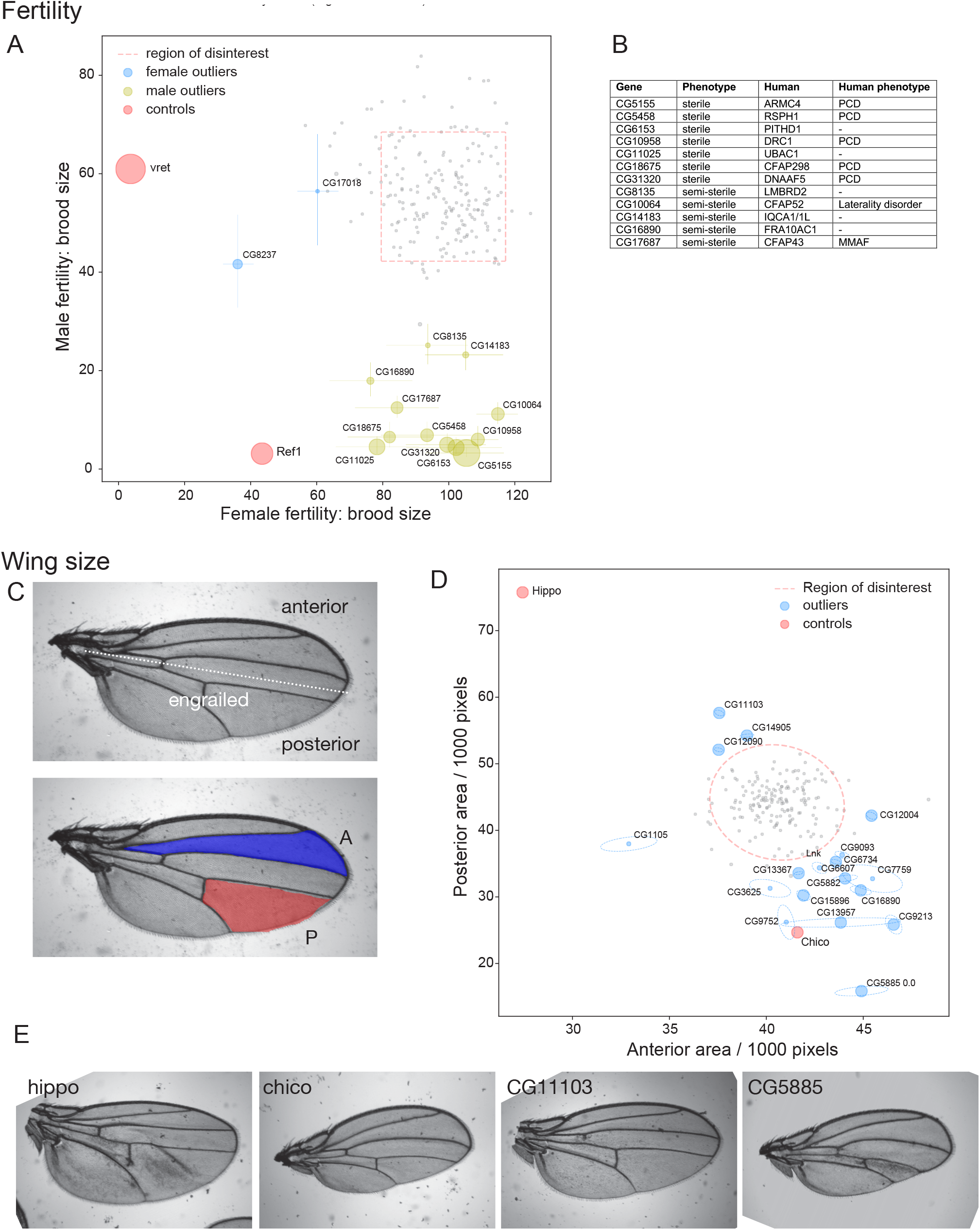
Testing of the unknome set of genes for roles in fertility and wing growth. (A) Plot of brood sizes obtained from matings in which each gene was knocked down in either the male or female germline. Dotted lines indicate outlier boundaries, with the genes named being those whose position outside of the boundary is statistically significant, error bars show standard deviation and the size of the circles is inversely proportional to the *p*-value (Tables S2 and S3). Controls shown are genes known to effect fertility in females (vret), or both males and females (Ref1). (B) Summary of the significant hits from the test of male fertility, showing the human ortholog and the phenotype reported for patients with loss of function mutations (PCD, primary ciliary dyskinesia; MMAF, multiple morphological abnormalities of the sperm flagella). (C) Adult wing illustrating the posterior domain that expresses engrailed during development and hence the engrailed-Gal4 driver used to express the hairpin RNAs. Also shown are the intervein areas measured to assess tissue growth in the anterior and posterior halves of the wing. (D) Plot of the mean area of the anterior and posterior intervein areas as in (C) for flies in which each gene was knocked down by RNAi in the posterior domain. Errors are shown as tilted ellipses with the major/minor axes being the square roots of the eigenvectors of the covariance matrix. Dotted lines indicate the outlier boundary, with the genes named being those whose position outside of the boundary is statistically significant, with the size of the circles being inversely proportional to the *p*-value (Tables S2 and S3). The genes Hippo (growth repressor) and Chico (growth stimulator) are included as controls. (E) Representative wings from flies expressing hairpin RNA for the indicated genes in the posterior domain. Hippo and Chico are controls as in (D), with CG11103 and CG5885 showing an increase or decrease in the posterior domain, respectively.

### - Female fertility

Two genes gave a partial, but significant, reduction in female brood size. During the course of our work, a mouse ortholog, MARF1, of one of these hits, CG17018, was identified in a genetic screen as being required for maintaining female fertility, apparently by controlling mRNA homeostasis in oocytes (Nishimura et al., 2018; Su et al., 2012; Yao et al., 2018). A recent study of CG17018 has confirmed that it is indeed required for female fertility in *Drosophila*, despite lacking some domains present in MARF1. Its appearance as a hit in our screen is therefore an encouraging validation of the approach (Zhu et al., 2018). The other gene, CG8237, has not previously been linked to fertility, but has a mammalian ortholog (FAM8A1) that has been recently proposed to help assemble the machinery for ER-associated degradation (ERAD) and so may have an indirect effect on oogenesis (Schulz et al., 2017; Zhu et al., 2017). We selected CG8237 for validation by CRISPR/Cas9 gene disruption as described below.

### - Male fertility

Seven genes showed near complete male sterility, with five further genes gave a reduction in brood size that was statistically significant. In humans, male sterility is one of the symptoms associated with primary ciliary dyskinesia (PCD), a disorder affecting motile cilia and flagella (Horani et al., 2016). Whilst our analysis was in progress, exome-sequencing allowed the identification of many new PCD genes (Horani and Ferkol, 2018; Legendre et al., 2021). Interestingly, five of the genes identified in our assay are homologs of human PCD genes (Figure 3B), of which CG5155 and CG31320 have since been shown to be required in *Drosophila* for male fertility (Cheng et al., 2013; Diggle et al., 2014). All of these genes comprise, or help assemble, the dynein-based system that drives the beating of cilia and flagella. In addition, human orthologs of two of the semi-sterile hits in the Unknome screen have been found to be mutated in related familial conditions. CFAP43 (orthologous to CG17687) is mutated in patients with multiple morphological abnormalities of the sperm flagella (MMAF), and CFAP52 (orthologous to CG10064) is mutated in laterality disorder, a condition caused by defects in ciliary beating during development (Coutton et al., 2018; Ta-Shma et al., 2015). A further semi-sterile hit, CG14183, is an ortholog of DRC11, a subunit of the nexin-dynein regulatory complex that regulates flagellar beating in *Chlamydomonas* (Gui et al., 2019). These findings prove the value of the Unknome database approach to identifying new genes of biological significance, and validate the RNAi-based screening approach.

Of the four remaining genes that showed male fertility defects, CG11025 is now only partially unknown as its human ortholog is a non-catalytic subunit of the Kip1 ubiquitination-promoting complex, an E3 ubiquitin ligase (Kravtsova-Ivantsiv et al., 2015). CG11025 was recently identified in a genetic screen for defects in ciliary traffic, and found to be required for fertility (Li et al., 2020). However, as described above, the unknome continues to erode only slowly, and the other three genes, CG8135, CG6153 and CG16890, remain poorly understood in any species. They are less likely to be flagellar components as they are not predominantly expressed in testes and, as described below, two were selected for validation by CRISPR/Cas9 gene disruption, along with CG10064 whose ortholog is mutated in laterality disorder.

### Contribution of unknome genes to tissue growth

To test the unknome set of genes for roles in tissue formation and growth we examined the effect of knocking down them in the posterior compartment of the wing imaginal disc, and then compared the area of the posterior compartment of the adult wing to that of the control anterior compartment (Figure 3C), a method previously used to detect effects of a range of different genes (Hahn et al., 2013; Ibar and Glavic, 2017). As controls, we used Hippo, a negative regulator of tissue size, and Chico, a component of the PI 3-kinase pathway that stimulates organ growth (Böhni et al., 1999; Irvine and Harvey, 2015). Knockdown of three of the unknome genes in the posterior compartment caused a statistically significant increase in its area (Figures 3D and 3E). These include CG12090, the *Drosophila* ortholog of mammalian DEPDC5, which was recently found to be part of the GATOR1 complex that inhibits the Tor pathway. Mutants in GATOR1 subunits promote cell growth by increasing Tor activity (Bar-Peled et al., 2013; Wei et al., 2016). The other two are CG14905 and CG11103. CG14905 is a paralog of a testes-specific gene CG17083, and both are orthologs of mammalian CCDC63/CCDC114 that have a role in attaching dynein to motile cilia, although CG14905 seems likely to have additional roles as it is ubiquitously expressed (Hjeij et al., 2014). CG11103 encodes a small membrane protein that shares a TM2 domain with Almondex, a protein with an uncharacterised role in Notch signalling (Michellod and Randsholt, 2008). We therefore selected CG11103 for further validation by CRISPR/Cas9 as described below.

A larger number of genes caused a reduced compartment size when knocked down (Figure 3D). However, this could arise from a wide range of causes and so this is broad ranging assay for protein importance, and indeed mammalian orthologs of several of the stronger hits have been subsequently found to act in known cellular processes such membrane traffic (CG13957, the ortholog of human WASHC4), lipid degradation (CG3625/AIG1) or tRNA production (CG15896/PRORP; CG9752/C9orf72). The strongest effect was seen with CG5885, an ortholog of a subunit of the translocon-associated protein (TRAP) complex that is associated with the Sec61 ER translocon (Russo, 2020). TRAP’s role is enigmatic and so it was also selected for CRISPR/Cas9 validation.

### Contribution of unknome genes to protein quality control

The removal of aberrant proteins is a fundamental aspect of cellular metabolism, and thereby organismal health, but it is a function that does not necessarily contribute substantially to well-screened developmental phenotypes. It also exemplifies our suspicion that a disproportionately high number of the unknome set of genes may be involved in quality control and stress response functions, which are likely to have been missed by many traditional experimental approaches. We therefore tested the unknome gene set for protein quality control phenotypes, using an assay based on aggregation of GFP-tagged polyglutamine, a structure found in mutants of huntingtin that cause Huntington’s disease (Zhang et al., 2010). When this Httex1-Q46-eGFP reporter is expressed in the eye, the aggregates can be detected by fluorescent imaging (Figure 4A). The RNAi guides were co-expressed in the eye to knock down unknome genes, and the number of polyQ aggregates quantified. Although there was considerable variation in aggregate number, statistical analysis allowed the identification of clear outliers among the unknome RNAi set. Most of the genes showing the largest increase in aggregates remain of unknown function (CG7785 (SPRYD7 in humans), CG16890 (FRA10AC1), CG14105 (TTC36), and CG18812 (GDAP2)), although mutation of GDAP2 in humans causes neurodegeneration, consistent with a role in quality control (Eidhof et al., 2018). More is now known about two of the hits. CG4050 is a mammalian ortholog of TMTC3, one of a family of ER proteins recently shown to be O-mannosyltransferases; deletion of TMTC3 causes neurological defects (Farhan et al., 2017; Li et al., 2018). CG5885 is the ortholog of the SSR3 subunit of the TRAP complex that also showed reduced wing size; in mammalian cells the TRAP complex is upregulated by ER stress (Russo, 2020). These hits are consistent with reports that ER stress can increase cytosolic protein aggregation (Hamdan et al., 2017). We selected CG5885 and CG16890 for CRISPR/Cas-9 validation, as described below.

**Figure 4.**
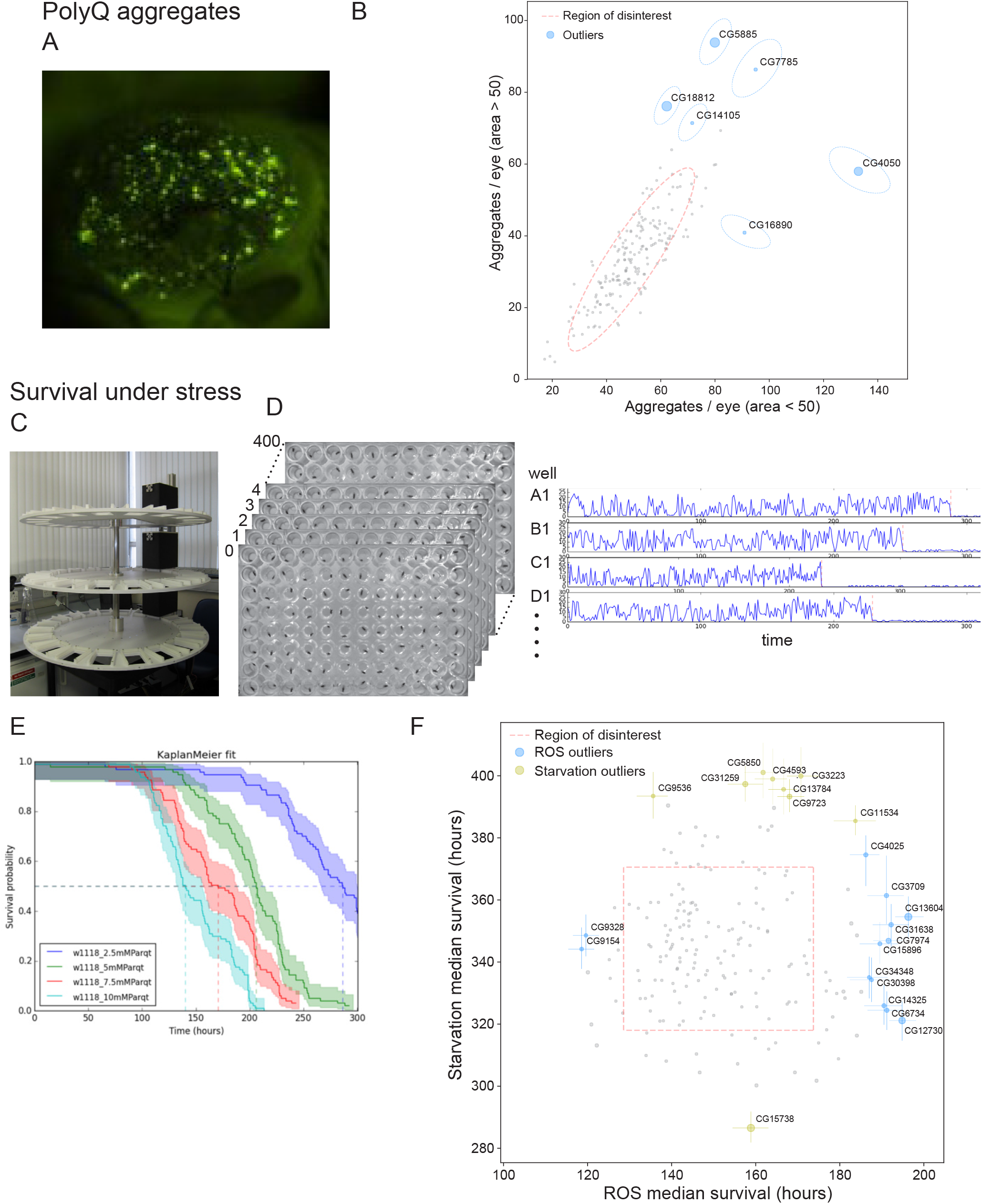
Testing of the unknome set of genes for roles in quality control and responses to stress. (A) Fluorescence micrograph of an eye from a representative stock expressing Httex1-Q46-eGFP under the control of the GMR-GAL4 driver. The GFP fusion protein forms aggregates whose number and size increase over time. (B) Plot of the mean number of large (>50 pixels) or small (<50 pixels) aggregates of Httex1-Q46-eGFP formed after 18 days in flies in which the unknome set of genes has been knocked-down by RNAi. Errors are shown as tilted ellipses with the major/minor axes being the square roots of the eigenvectors of the covariance matrix. Dotted lines indicate an outlier boundary set at 90% of the variation in the dataset, with the genes named being those whose position outside of the boundary is statistically significant with a p-value <0.05, with the size of the circles being inversely proportional to the *p*-value (Tables S2 and S3). (C) Flywheel system apparatus for time-lapse imaging of 96 well plates containing one fly per well. (D) Use of time-lapse imaging to assay viability. 96 well plates were imaged very hour and the movement between frames quantified for the fly in each well. Plots of movement size over time allow time point for cessation of movement and hence loss of viability to be determined automatically. (E) Survival plots obtained with the flywheel for flies on food containing the indicated concentration of oxidative stressor paraquat. Increased levels of the paraquat shorten survival times. (F) Plot of the mean survival time of fly lines in which the unknome set of genes has been knocked-down by RNAi and which were then exposed to paraquat to induce oxidative stress or were starved for amino acids. Dotted lines indicate an outlier boundary set at 80% of the variation in the dataset, with the genes named being those whose position outside of the boundary is statistically significant (*p-*value <0.05), with error bars showing standard deviation and the size of the circles inversely proportional to the *p*-value (Tables S2 and S3).

### Contribution of unknome genes to resilience to stress

Genomes have evolved to deal with many environmental stresses, and again these are processes poorly investigated by traditional genetic approaches. We therefore tested resilience to stress, following knock down of the unknome set. To quantify the viability of large numbers of flies, individual flies were arrayed in 96 well plates, and the plates maintained on a “flywheel” that rotated them under a camera every hour (Figure 4C). Viability was indicated by movement between images, allowing time of death to be determined with an accuracy of +/- one hour (Figures 4D and 4E). We applied this method with two challenges likely to be associated with different cellular resilience mechanisms: amino acid starvation and oxidative stress.

### - Resilience under starvation

Under conditions of amino acid deprivation, knock down of eight of the unknome test set significantly prolonged survival (Figure 4F). Seven of these genes remain of unknown function, but interestingly, five have orthologs in other species whose localisation or interactions suggest that they have roles in the endosomal system. Thus DEF8, the mammalian ortholog of CG11534, has been reported to interact with Rab7 (Fujiwara et al., 2016; Gillingham et al., 2014), and TMEM184A (CG5850) has been reported to act in the endocytosis of heparin (Pugh et al., 2016). In addition, the mammalian orthologs of CG4593 and CG9536 (CCDC25 and TMEM115) are Golgi localized proteins of unknown function, and the yeast ortholog of CG13784 (ANY1) has been found to suppress loss of lipid flippases that act in endosome-to-Golgi recycling (Ong et al., 2014; Takar et al., 2019). Our identification of this cluster of genes with related functions suggests that defects in endocytic recycling can prolong survival in starvation, possibly by altering autophagy or by reducing signalling from receptors that promote anabolism. The other two genes that improved starvation resilience when knocked down have no known function in any species, with loss of CG31259 (TMEM135) causing mitochondrial defects, and nothing reported for CG3223 (UBL7) (Lee et al., 2016; Shibano et al., 2015). One gene, CG15738, caused an increased susceptibility to starvation, and it has been found to be an assembly factor for mitochondrial complex I, whose loss compromises viability (Zhang et al., 2013).

### - Resilience under oxidative stress

Resistance to oxidative stress was tested with paraquat, an insecticide widely used to elevate superoxide levels in *Drosophila* (Phillips et al., 1989; Rzezniczak et al., 2011). There was considerable variability in the survival times, but eleven genes gave a statistically meaningful increase in resistance (Figure 4F). Most of these genes remain unknown, but three have since been reported to have functions related to oxidative stress signalling. The mammalian ortholog of CG4025 (DRAM1/2) is induced by p53 in response to DNA damage and promotes apoptosis and autophagy (Guan et al., 2015). The mammalian orthologs of CG13604 (UBASH3A/B) are tyrosine phosphatases that repress SYK kinase, an enzyme reported to help protect cells against ROS, with superoxide activation of *Drosophila* Syk kinase signalling tissue injury (Secchi et al., 2015; Srinivasan et al., 2016; Tsygankov, 2018). Finally, the ortholog of CG3709 in archaea has tRNA pseudouridine synthase activity, but the human ortholog PUS10 has been reported to be cleaved during apoptosis and promote caspase-3 activity, thus its loss may slow apopotic cell death (Jana et al., 2017). Of the other eight hits, five remain poorly characterised, one is involved in mitochondrial function and so may reduce ROS production, and two are involved microtubule function with no clear link to superoxide responses. Although further validation will be required, these five genes seem good candidates to have a role in mitochondria or ROS-response pathways.

### Contribution of unknome genes to locomotion

Metazoans benefit from a musculature under neuronal control. We therefore addressed the possibility of neuromuscular functions by testing the role of the unknome set of genes in locomotion, using the iFly tracking system, in which the climbing trajectories of adult flies are quantified by imaging and automated analysis (Figure 5A) (Jahn et al., 2011; Kohlhoff et al., 2011). Climbing speed declines with age, so the assay was performed at both 8 days and 22 days post eclosion. Climbing speeds are inevitably somewhat variable, even in wild-type flies, but nonetheless six genes were statistically significant outliers when assayed after 8 days (Figure 5B). Two of these genes remain poorly understood, and for three of the others recent work indicates a role in muscle or neuronal function. These include CG9951, whose human homolog CDCC22 has been recently found to be a subunit of the retriever complex that acts in endosomal transport. Missense mutations in CDCC22 causing intellectual disability (McNally et al., 2017; Voineagu et al., 2012). The human ortholog of CG13920 (TMEM35A) is required for assembly of acetylcholine receptors (Matta et al., 2017). Finally, CG3479 is the gene mutated in the *Drosophila outspread (osp)* wing morphology allele, and is expressed in muscle; one of its two mammalian orthologs MPRIP has been found to regulate actinomyosin filaments (McNabb et al., 1996; Surks et al., 2005).

**Figure 5.**
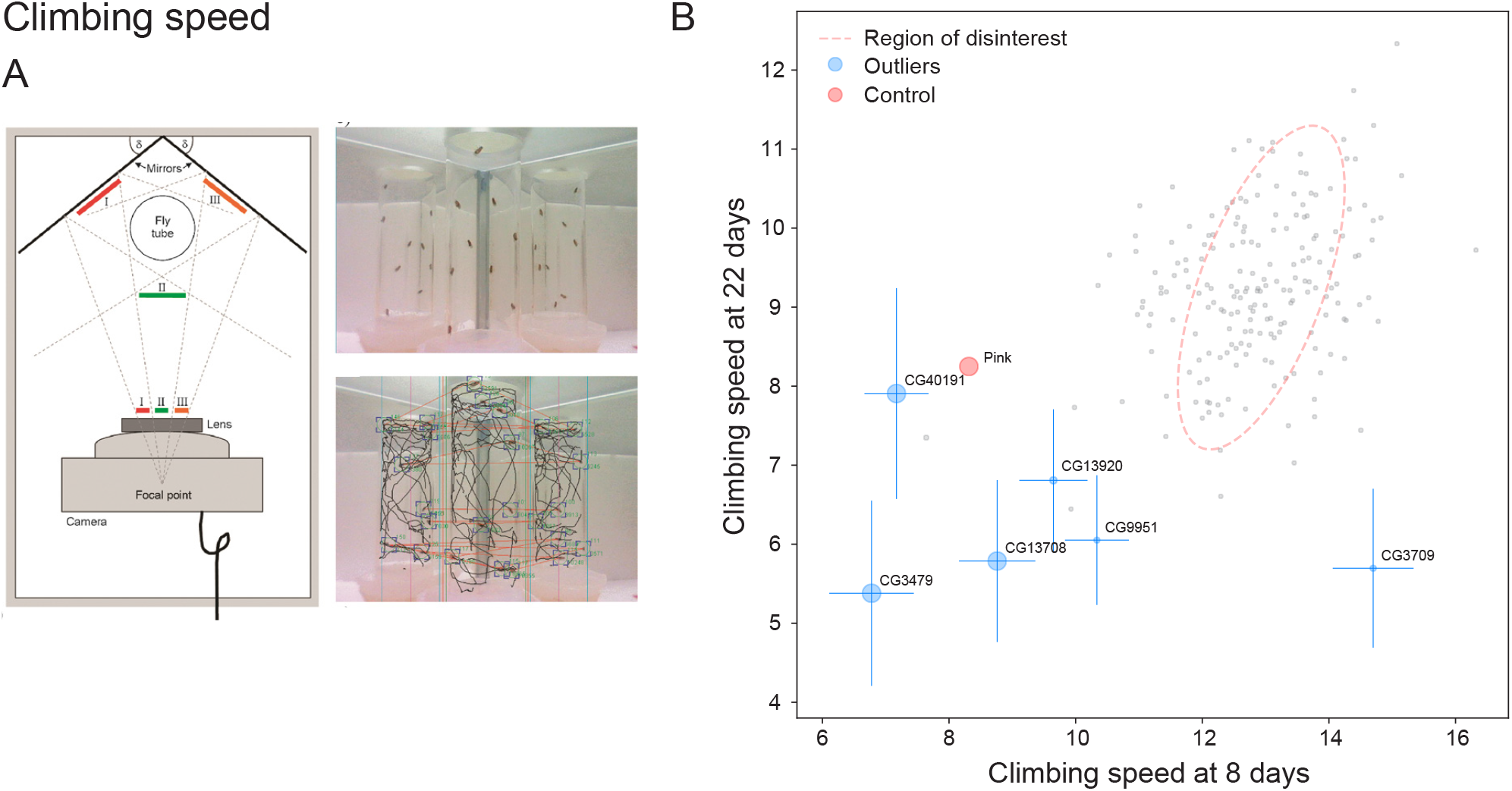
Testing the unknome set of genes for roles in locomotion. (A) iFly tracking system for automatic quantitation of *Drosophila* locomotion (reproduced from Kohlhoff et al (2011) (Kohlhoff et al., 2011)). (B) Plot of the mean climbing speeds of fly lines in which the unknome set of genes has been knocked-down by RNAi, and the speeds for each line were determined after eight days or 22 days post eclosion. The Parkinson’s gene Pink1 was included as a positive control (Clark et al., 2006). Dotted lines indicate an outlier boundary set at 90% of the variation in the dataset, with the genes named being those whose position outside of the boundary is statistically significant with a p-value <0.1, with error bars showing standard deviation and the size of the circles inversely proportional to the *p*-value (Tables S2 and S3).

### Validation of fertility screen hits by gene disruption

Analysis of gene function by RNAi can be confounded by off-target effects. We therefore used CRISPR/Cas9 gene disruption to validate selected hits from two of the phenotypic screens. From the fertility screens, three male steriles and one female sterile were selected for genetic disruption. Of the male hits, CG10064 and CG6153 were both confirmed as being required for male fertility (Figures 6A to 6D). CG10064 is a WD repeat protein, and mutation of its human ortholog, CFAP52, results in abnormal left-right asymmetry patterning, a process known to depend on motile cilia (Ta-Shma et al., 2015). Male flies lacking CG10064 produced motile sperm, but following mating they did appear to not persist in the female’s sperm storage organ, the seminal receptacle, indicating that the protein is required for the optimal function rather than formation of motile cilia (Bloch Qazi et al., 2003). CG6153 comprises a PITH domain that is also found in TXNL1, a thioredoxin-like protein that associates with the 19S regulatory domain of the proteasome through its PITH domain (Andersen et al., 2009; Wiseman et al., 2009). Males lacking CG6153 made morphologically normal sperm, but they did not accumulate in the seminal vesicle, the organ in which nascent sperm are stored prior to deployment, suggesting that they have limited viability (Figures 6E to 6J). Neither CG6153, nor its human ortholog PITHD1, are testis specific, and indeed orthologs are also present in non-ciliated plants and yeasts, suggesting that the protein has a role in an aspect of proteasome biology that is of particular importance for maturing viable sperm. Recent work on mouse PITHD1 indicates it has a role in both olfaction and fertility (Kondo et al., 2020; Lachén-Montes et al., 2020). The other male sterile hit, CG16890, and the female sterile hit CG8237 did not show reduced fertility when disrupted and presumably represent off-target RNAi effects (fig. S1).

**Figure 6.**
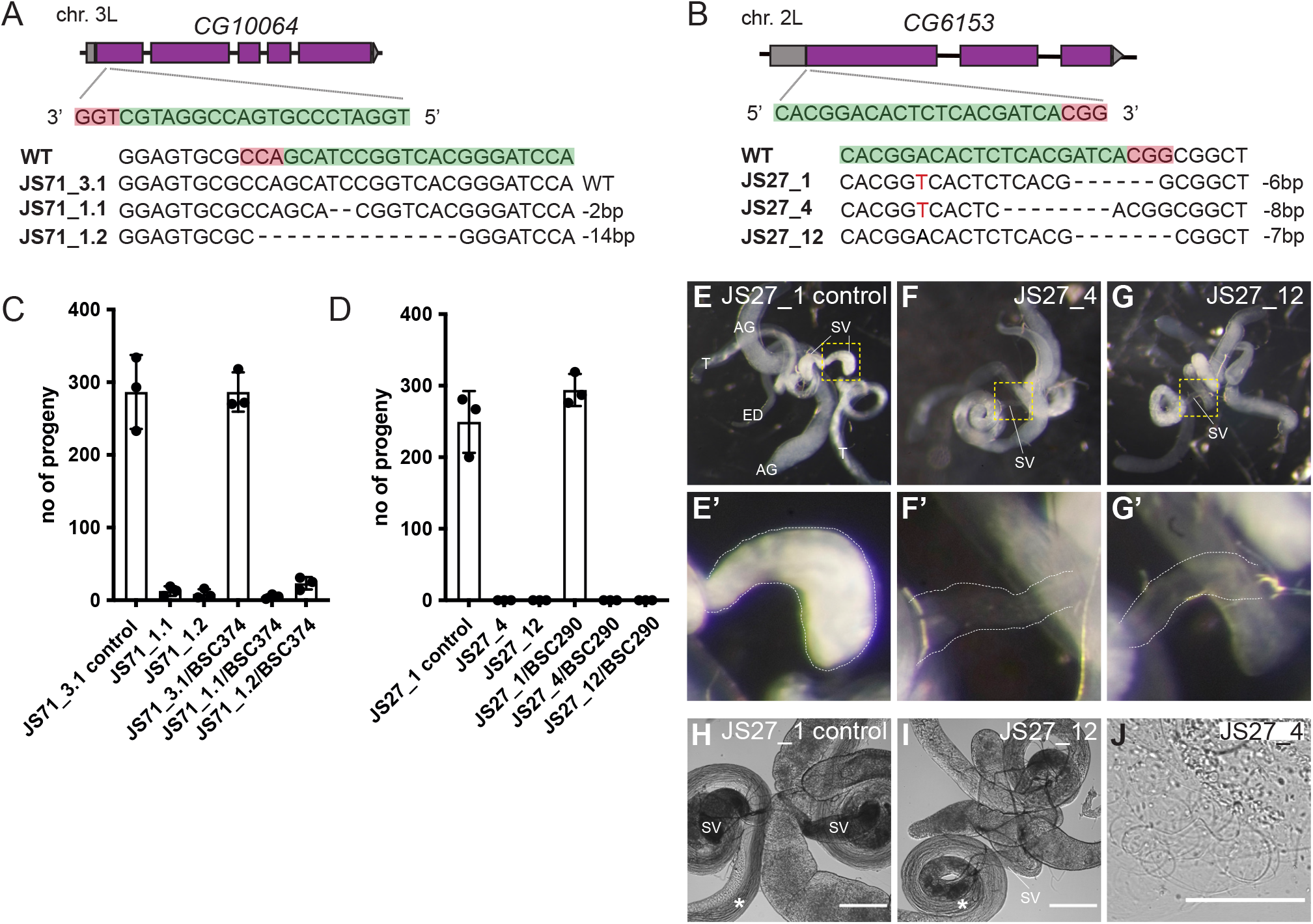
Validation of RNAi male sterility phenotypes using CRISPR/Cas9 gene disruption. (A,B) Schematics of the genomic locus of candidate genes, position of CRISPR target sites and mutant alleles analysed. (C,D) Assessment of male fertility of mutants (homozygous and over a deficiency). The graphs show mean values +/-SD of the number of progeny produced by mutant males. Three crosses with 5 wild-type virgins and 3 mutant males were analysed for each genotype. Wild-type males and/or males carrying in-frame mutations were used as controls. Where possible, alleles covering both alternative reading frames were analysed. (E-G) Widefield fluorescent micrographs of male reproductive systems of control and JS27/CG6153 mutants expressing Don Juan-GFP to label sperm (F-G). Mutants exhibit empty seminal vesicles, (E’-G’) showing zoomed regions of seminal vesicles from E-G (yellow dashed squares). (H-J) Widefield phase micrographs of reproductive systems of control and mutant males. Sperm are produced in both (asterisks), suggesting that sperm are made in the mutant but do not survive. Note that some mutant sperm gets into the ejaculatory duct (J). AG: accessory gland, ED: ejaculatory duct, SV: seminal vesicle, T: testis. Scale bars, 200 µm (H-I), 100 µm (J).

### Wing size hit CG11103 is a regulator of Notch signalling

Knockdown of gene CG11103 caused alterations in the growth of the wing (Figures 4D and 4E). When CG11103 was removed with CRISPR/Cas9, mutant females and males were viable without any obvious phenotypes, but females were completely sterile. Eggs laid by mutant females were fertilised but failed to develop (Figures 7A to 7D). Cuticle preparations and antibody labelling of the pan-neuronal marker Elav showed a hyperplasia of nervous system at the expense of the epidermis (Figures 7F and 7G). This phenotype is characteristic of defects in the highly conserved Notch signalling pathway which is required in the *Drosophila* embryo to determine/specify the neuroblasts that give rise to the CNS in a process called lateral inhibition. CG11103 contains a TM2 domain that comprises two transmembrane domains connected by a short linker (Kajkowski et al., 2001). The function of this domain is unknown, but it occurs in two related proteins in *Drosophila*, and all three of the fly proteins have human orthologs (Figure 7B). Interestingly, one of these, *almondex*/*CG12127*, was identified as a gene required for Notch signalling in embryos, although its role remains unclear (Michellod et al., 2003). The third related gene, *CG10795*, is also of unknown function, so we knocked it out with CRISPR/Cas-9 and discovered that it too showed phenotypes indicative of a severe defect in Notch signalling (Figures 7H to 7L). Thus, all three proteins are required for a cellular process essential for embryonic Notch function. All three human TM2D proteins were hits in a recent genome-wide screen for defects in endosomal function (Haney et al., 2018), and endosomes play a critical role in Notch signalling. Further work will be required to determine the precise role of these proteins, and how it relates to wing growth, but their likely role in endosomal function, combined with the existence of related TM2 domain proteins in bacteria and archaea, suggest fundamental role in cell function rather than an exclusive role in Notch signalling.

**Figure 7.**
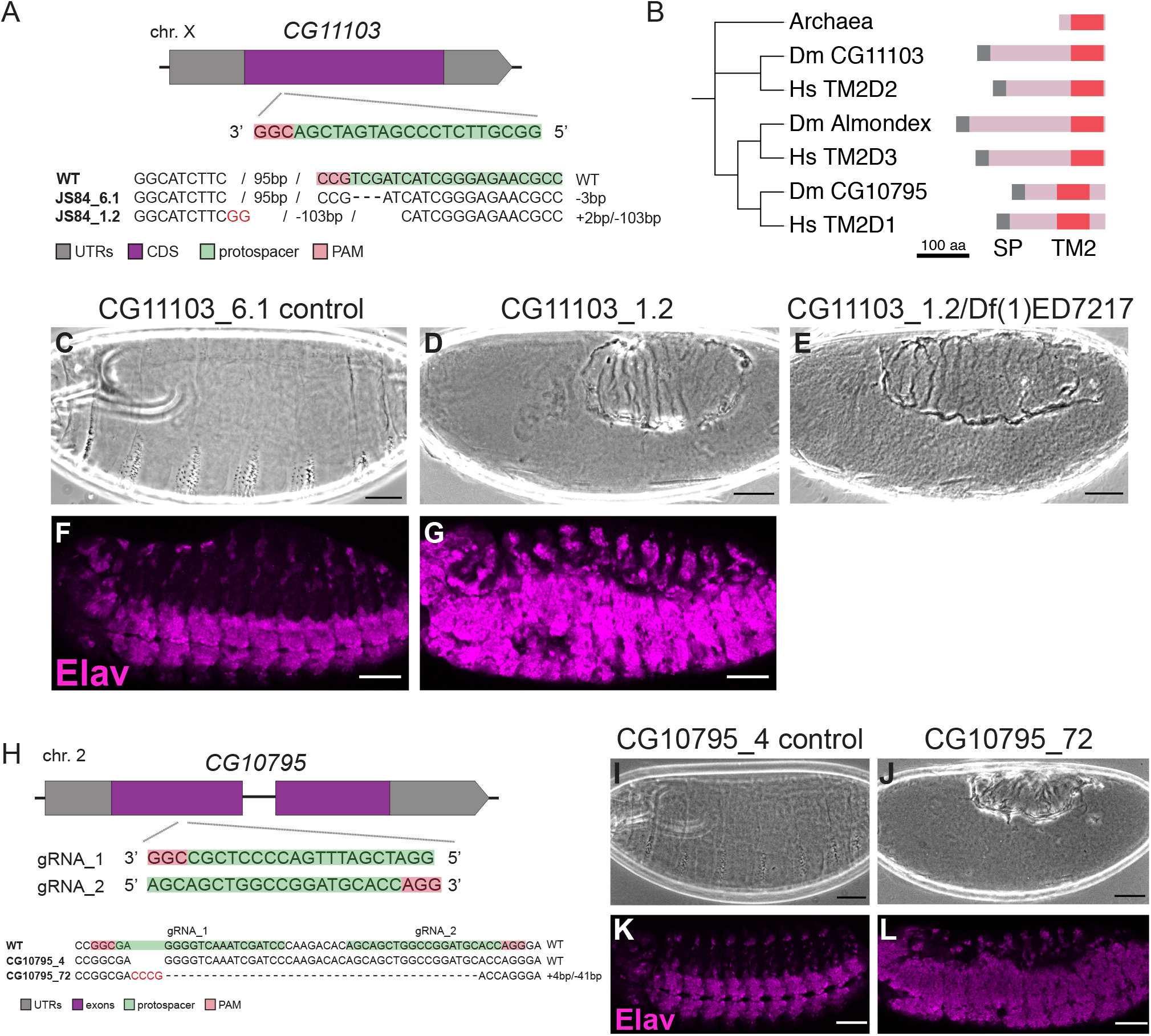
Investigation of wing growth hit *CG11103* using CRISPR/Cas9 gene disruption. (A) Schematic of the genomic locus of candidate *CG11103*, position of the CRISPR target site and the mutant allele analysed. Flies carrying an in-frame mutation were used as control. (B) Gene tree for TM2 domain proteins in humans and *Drosophila*, with an archaeal TM2 protein as an outlier. Tree built using sequence of TM2 domains alone using T-Coffee. A fourth TM2 domain protein is present in *Drosophila* and humans, Wurst/DNAJC22 which has additional TMDs and a DNAJ domain and appears to play a role in clathrin-mediated endocytosis (Behr et al., 2007). (C-E) Cuticle phenotypes of embryos laid by control females and mutant females (homozygous or over a deficiency). (F,G) Micrographs of embryos laid by control females and homozygous mutant females stained against the pan-neuronal marker Elav. Scale bars: 50 µm. (H) Schematic of the genomic locus of *CG10795*, position of CRISPR target sites and the alleles analysed. Flies without an indel were used as control (*CG10795_4*). (I,J) Cuticle phenotypes of embryos laid by control or mutant females. (K,L) Micrographs of embryos laid by control or mutant females stained for the pan-neuronal marker Elav. Scale bars: 50 µm.

Taken together, this genetic validation data confirms that the RNAi screening approach, despite its known caveats, has given accurate phenotypic information for at least a substantial subset of the hits from our RNAi screens of the unknome set of genes.

## Discussion

The totality of scientific knowledge represents the summed activity of numerous individual research groups, each focusing on specific questions whose selection is influenced by many factors, some scientific and some more socially determined. The latter class include issues like a preference for the relative safety, sociability and kudos available when working in well-established fields, but is also strongly influenced by funding mechanisms. These usually aim to address societal needs but are subject to subjective assessment, historical precedent and political pressures. In particular, the need to justify proposed research with reference to an established body of work, and preliminary data, can restrict investigation into truly unknown areas. Putting it more positively, there is potential for scientific progress to be accelerated disproportionately by identifying situations where questions are being inadvertently and unjustifiably neglected. We have directly addressed here an area of long-standing concern: that biological research largely ignores less well known, but potentially important, genes (Edwards et al., 2011; Oprea et al., 2018; Sinha et al., 2018; Stoeger et al., 2018). Our results support the conclusion that this concern is well founded.

Our approach has been to develop an Unknome database. This has confirmed previous observations that poorly understood genes are relatively neglected; we also find that this problem is persisting even though there has been some progress in assigning functions to some of these genes. Recent developments in exome sequencing have allowed the identification of novel components of pathways whose genes give a well-defined set of disease symptoms, as has been seen with the cilia proteins identified from patients with ciliopathies (Horani et al., 2016; Legendre et al., 2021). In addition, the advent of the CRISPR/Cas9 system has enabled screens that cover whole genomes (*13, 90, 91*). However, such screens are typically performed in cultured cells and hence cover only a subset of biological processes, and can also miss genes that have closely related, and thus functionally redundant, paralogs (De Kegel and Ryan, 2019).

We used the Unknome database to select about 200 genes that appeared both highly conserved and particularly poorly understood, and then applied functional assays in whole animals that would be impractical at genome-wide scale. Using seven assays, designed to interrogate defects in a broad range of biological functions, we found phenotypes for 59 genes, in addition to the 62 genes we had found to be essential for viability (Table S4). Our approach relied on RNAi, but when seven of the hits (corresponding to six genes) were retested with CRISPR/Cas9 gene disruption we could validate four, suggesting a true hit rate of ∼50%, a good outcome for an RNAi-based approach. The use of RNAi to knock down candidate genes is powerful in this context because it allows for tissue specific knock down; moreover, the likely incomplete loss of function achieved by RNAi can allow essential genes to reveal otherwise hidden hypomorphic phenotypes. Conversely, we note that as CRISPR approaches become ever more streamlined and sophisticated, future exploitation of the Unknome database can realistically use CRISPR technology to investigate functions of unknown genes.

An important primary conclusion of our work is that these uncharacterised genes have not deserved their previous neglect, a conclusion strengthened by a variety of other studies published during the course of our studies, again revealing important functions for unknown genes. Again, this highlights the gradual shrinking, albeit slowly, of the unknome. Perhaps, most significantly, our database provides a powerful, versatile and efficient platform to identify and select important genes of unknown function for analysis, thereby accelerating the closure of the gap in biological knowledge that the unknome represents. In practical terms, the Unknome database provides a resource for researchers who wish to exploit the disproportionate opportunities associated with unstudied areas of biology.

Thinking about how to evaluate ignorance of gene function guided our bioinformatic approach that allowed selection of a set of genes small enough for complex phenotypic screening in a whole animal. At a broader level, we believe that acknowledging and evaluating ignorance is an important factor in decisions about the relative priority given to addressing the remaining fundamental questions in biology, versus translating and exploiting what we already know. However, ignorance can only have value if it can be meaningfully measured. Developing the Unknome database highlighted a couple of issues that affect our assessment of the state of knowledge of gene function. First, our approach relied on identifying orthologs from major organisms used for biological research. Although current methods for ortholog identification work well, there is still scope for improvement (Altenhoff et al., 2016; Glover et al., 2019; Hu et al., 2011; Wang et al., 2017). Our approach also relied on the comprehensive and systematic annotation of gene function by the Gene Ontology (GO) Consortium (Ashburner et al., 2000; The Gene Ontology Consortium, 2019).

A second issue that arises from our work is that the current rapid rate of genome sequencing has required that most annotation is now automated rather than manual. This has led to the development of powerful methods to add functional annotation based on similarities to genes from other species (Schnoes et al., 2013). However, such methods aim to cumulatively add annotation rather than remove disproven conclusions or address contradictions, as this requires time-consuming manual curation. Moreover, increasing numbers of functional annotations are based on phenotypes from high-throughput screens for genetic phenotypes or protein-protein interactions, both of which are prone to generating false positives (Schnoes et al., 2013). Thus, genes inevitably accrete annotations over time, some of which may be wrong, contradictory or superficial but have little prospect of being corrected in the foreseeable future. As a result, the admirable aim of adding new gene annotation carries the risk of inadvertently obscuring our understanding of what is genuinely unknown.

An illustration of this problem is the gene CG9536 (TMEM115 in humans). This protein has been annotated as having endopeptidase activity based on distant sequence similarity to the rhomboid family of intramembrane proteases. However, CG9536, and its relatives in other species, lack the conserved residues that form the active site in rhomboids, and thus the only thing that can be currently concluded about the function of CG9536 is that it is almost certainly *not* a protease (Freeman, 2014). A more extreme case is htt, the *Drosophila* ortholog of huntingtin. This was not in the unknome test set because the extensive study of the role of huntingtin in human disease has led to many preliminary suggestions of function which have resulted in annotations linked to transcription, transport, autophagy, mitochondrial function etc, and yet the current consensus is that huntingtin’s precise cellular role remains uncertain (Saudou and Humbert, 2016).

In conclusion, we find that accurately evaluating ignorance about gene function provides a valuable resource for guiding biological studies, and may even be important for determining strategies to efficiently fund science. We have developed an approach to tackle directly the huge but under-discussed issue of the large number of well conserved genes that have no reliably known function, despite the likelihood that they participate in major and even possibly completely new areas of biological function. We hope that our work will inspire others to define and characterise further the unknome, and also to seek means to ensure that gene annotation accurately preserves and recognises our current state of ignorance.

## Methods

Please see Supplemental Information (below).

## Acknowledgements

We thank Damian Crowther for loan of the iFly tracking system, Sara Imarisio for advice on proteostasis assays, the LMB workshops for help with the system for lifespan measurements, Anna Parish for fly stock maintenance, and Manu Hegde for comments on the manuscript. Funding was from the MRC (MRC file reference number MC_U105178783). Funding was from the Medical Research Council, as part of United Kingdom Research and Innovation (also known as UK Research and Innovation), file reference number MC_U105178783.

## Author contributions

JR: investigation, data curation, methodology; SAJ: investigation; TS: methodology, software; NM: investigation; RDJ: formal analysis; SE: investigation; CR: investigation; MF conceptualization, methodology, project administration; SM: conceptualization, methodology, project administration, writing.

## Supplemental Information

**Figure S1 Testing of RNAi sterility hits using CRISPR/Cas9 gene disruption**

**STAR Methods**

**Statistical Supplement**

### Supplemental Tables

**Table S1. Genes selected for unknome screen, related to Figures 3, 4, and 5** List of *Drosophila* genes used for the unknome screen and the corresponding RNAi stocks.

**Table S2. Data points from screens, related to Figures 3, 4, and 5**

Data for individual flies from batches assayed for each genotype in functional screens.

**Table S3. Mean and variances from screens, related to Figures 3, 4, and 5** Statistical analysis of batches assayed for each genotype in functional screens, as used for plots in figures.

**Table S4. Summary of hits from the screens, related to Figures 3, 4, and 5** Genes whose knockdown gave significant effects in the functional screens.

### Supplemental Movie

Movie S1. **Lifespan assay in 96 well plate, related to Figure 4**

Representative time-lapse movie of flies in a 96 well plate. Frames captured every hour and played at 30 frames/second.

**Figure S1.**
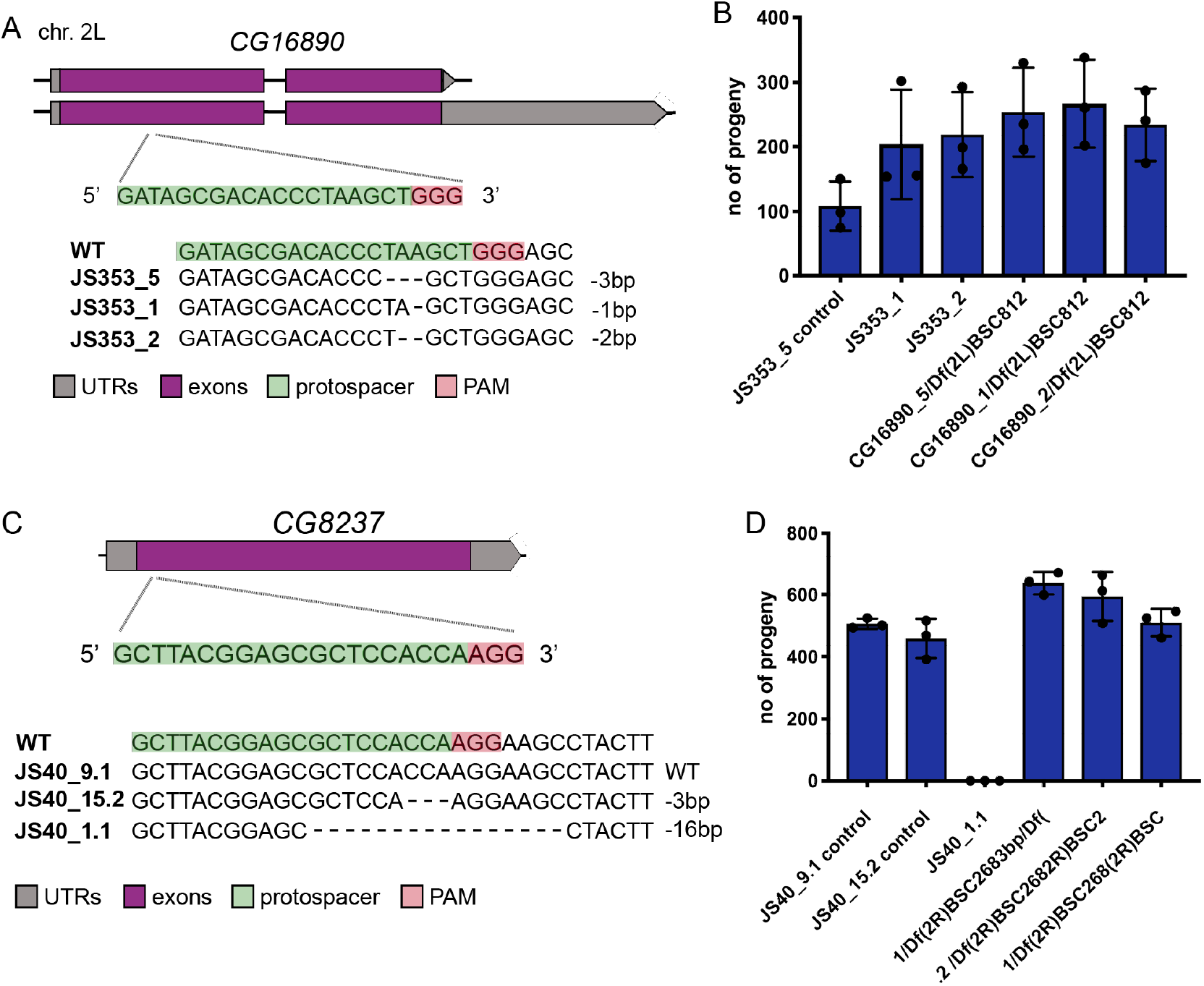
Testing of RNAi sterility hits using CRISPR/Cas9 gene disruption. (A) Schematics of the genomic locus of candidate JS353/CG16890, position of CRISPR target sites and mutant alleles analysed. (B) Assessment of male fertility of mutants (homozygous and over a deficiency). The graphs show mean values +/-SD of the number of progeny produced by mutant males. Three crosses with 5 WT virgins and 3 mutant males were analysed for each genotype. WT males and/or males carrying in-frame mutations were used as controls. Alleles covering both alternative reading frames were analysed. (C) Schematic of the genomic locus of candidate JS40/CG8237, position of the CRISPR target site and the mutant allele analysed. (D) Assessment of female fertility of mutants (homozygous and over a deficiency). The graph shows mean values +/-SD of the number of progeny produced by mutant females. Three crosses with 5 mutant virgins and 3 WT males were analysed. WT males and males carrying an in-frame mutation were used as controls.

## STAR Methods

### Lead contact

Further information and requests for resources should be directed to and will be fulfilled by the lead contacts, Matthew Freeman and Sean Munro.

### Materials availability

Fly stocks generated during this study are available by request to the Lead Contact.

### Data and code availability

Data from the functional screens are available in the main text or the supplemental tables. Code for functional assays are available at https://github.com/tjs23/unknome.

### Experimental Model and Subjects Details

#### Drosophila

Hairpin RNAi stocks for the Unknome set were from the KK library of the Vienna Drosophila Resource Centre (Table S1). During the course of our studies it was reported that the stocks in this library have the transgene in one of two sites in the genome (the annotated locus 40D or the non-annotated site 30B), and insertions at 40D can cause lethality when the guide RNA is expressed (Green et al., 2014; Vissers et al., 2016). PCR analysis with the previously used diagnostic primers was applied to 360 of the 365 lines, with the five remaining lines being lethal when expressed and so not included in any of the functional screens. This PCR analysis revealed that 98 of the 360 lines have the transgene in the problematic 40D site, a frequency of 27%, comparable to the 23% (9/39) and 25% (38/150) found previously. All but one of these 98 lines gave a lethal or semi-lethal phenotype when crossed to the ubiquitous da-GAL4 driver (Table S1).

## Method Details

### Construction of the Unknome database

The of protein sequence data that we considered corresponds to the reference UniProt Proteomes [https://www.uniprot.org/proteomes/] used by the latest PantherDB and includes human and 11 model organism species: *A. thaliana, C. elegans, D. rerio. D. discoideum, D. melanogaster, E. coli* (K12), *G. gallus, M. musculus, R. norvegicus, S. cerevisiae* and *S. pombe* (Mi et al., 2021; UniProt Consortium, 2021).

The Unknome database aggregates relevant information from the listed sources and provides a default knownness score for each protein and protein family (cluster) and can be (re-)compiled in a few hours. Here PantherDB provides the protein family information, via a group of UniProt IDs, that can be combined with selected information from UniProt entries; including protein sequence, GO terms, PubMed citations, species, gene name(s) and cross-references to species-specific databases.

The GO terms present in each UniProt entry are automatically provided by the Gene Ontology Annotation (GOA) database [https://www.ebi.ac.uk/GOA] (Gene Ontology Consortium, 2021). Evidence terms from the OBO Foundry are employed by GO (Smith et al., 2007), and in the Unknome database they were weighted according to their evidence codes using the following default values: EXP; 0.8, IDA; 0.8, IPI; 0.8, IMP; 0.8, IGI; 0.8, IEP; 0.8, ISS; 0.5, ISO; 0.5, ISA; 0.5, ISM; 0.5, IGC; 0.3, RCA; 0.6, TAS; 0.9, NAS; 0.6, IC; 1.0, ND; 0.0, IEA; 0.0, NR; 0.0, IRD; 0.0, IKR; 0.0, IBA; 0.5, IBD; 0.5. (see http://geneontology.org/docs/guide-go-evidence-codes/ for a full description). After weighting they were summed to generate a knownness score for each protein. The knownness score for the family, as defined by PantherDB, was then the maximum score from all the protein members present in the human and model organism list.

All protein GO terms linked in the database were dated according to when they were first linked with the UniProt entry, so as to be able to track the historical change of knownness. Though this information is not directly accessible within UniProt entries, the GOA database makes this information available via GAF format files at ftp://ftp.ebi.ac.uk/pub/databases/GO/goa/UNIPROT/.

The Unknome database is currently available as SQLite Version 3 files and is presented with a web interface at the URL https://unknome.mrc-lmb.cam.ac.uk/. This website is constructed using the Python module Django and provides views on the underlying database with easy filtering by knownness. In particular, the site displays the change over time in knownness for each protein family, and lists all of the associated dated GO terms. The web site also makes all data available for download; from individual protein sequences to the whole SQL database file.

### *Drosophila* genetics

Expression of the RNAi hairpins was driven with either the ubiquitous driver da-GAL4 driver, or with tissue-specific drivers: en-GAL4 (wing), bam-GAL4-VP16 (male fertility), MTD-GAL4 (female fertility), and GMR-GAL4 (proteostasis in the eye). UAS-Dicer-2 was included in all cases except for the two fertility screens as this has been found to improve the efficiency of RNAi (Dietzl et al., 2007). For the proteostasis screen, the driver line also contained UAS-Httex1-Q46-eGFP (Zhang et al., 2010). In the lethality screen, those crosses that produced no adult progeny were defined as ‘lethal’, whilst those where the progeny reached the pharate stage but the majority could not hatch, and those that did failed to expand wings and did not survive were “semi-lethal”.

For validation using CRISPR/Cas9, the following fly stocks were used: nos-phiC3; attP40 (DBSC #25709), nos-phiC3;;attP2 (DBSC #25710), CFD2 (Port et al., 2014), TH_attP2 (Port et al., 2015), Df(1)ED7217 (DBSC #8952), Df(2R)BSC268 (DBSC #26501), Df(2L)BSC812 (DBSC #27383), Df(2L)BSC290 (BDSC #23675), Df(3L)BSC374 (BDSC #24398). Spermatids and sperm were labelled with Don Juan (dj)-GFP (Santel et al., 1997).

### Fertility

Fertility was monitored using competitive assays, in which one red-eyed fly expressing the RNAi and one white-eyed w1118 wild-type fly were placed with 4 wild-type flies of the opposite sex. For male fertility the Bam-Gal4 driver was used in combination with Dicer, and for female fertility MTD-Gal4 was used without Dicer. The flies were allowed to mate for seven days, transferring to fresh vials every 2-3 days. After seven days, the parental generation was removed and all progeny that emerged from the vial were counted, with eye colour used to determine the parent of each. Flies from the RNAi parent were separated, imaged, and quantified using Fiji image analysis platform (Schindelin et al., 2012), with a custom macro (https://github.com/tjs23/unknome). Individual data for both males and females is in Table S2, with the means and the variances errors plotted in the figures provided in Table S3.

### Wing growth assay

The genes in the unknome set were knocked down in the posterior half of the wing by using an engrailed-GAL4 driver combined with UAS-dcr-2. For each cross, at least 10 independent wings were collected, and mounted on a slide under a coverslip in 50% glycerol/PBST. Images obtained with a 5 x objective were analysed using a Fiji macro to contrast the veins from the rest of the wing (https://github.com/tjs23/unknome), and then the areas of specific inter-vein regions in the anterior and posterior halves were determined. Individual data is in Table S2, with the means and the variances errors plotted in the figures provided in Table S3.

### Proteostasis assay in the eye

To interrogate the handling of misfolded proteins, a GFP fusion to part of huntingtin with a polyglutamine repeat was expressed in eyes, and the number of GFP-positive aggregates determined (Zhang et al., 2010). UAS-Httex1-Q46-eGFP was expressed in the eye along with the RNAi using GMR-Gal4. One eye from each of nine males per genotype were imaged after 18 days at 25°C, using 3 males per independent cross. GFP-positive aggregates were quantified with Fiji using a custom macro that determined the area of the eye, and then scored aggregates that were either smaller or bigger that 50 pixels (https://github.com/tjs23/unknome). Individual data is in Table S2, with the means and the variances errors plotted in the figures provided in Table S3.

### Survival under stress

To measure lifespan under stress we developed an automated system for following viability over may days. Flies were placed 96 well plates and photographed every hour with image analysis then used to identify when the flies stopped moving. To prepare the plates, low nitrogen-free fly food was placed at the bottom of each well (8 g agar, 50 g glucose, and 5 g pectin per litre with 0.25% nipagin, antibiotics, and 4 ml/litre propionic acid as preservative). To assay oxidative stress, the same food was used with the addition of 7.5 mM paraquat. Adult male flies were subdued with CO2 and single flies placed in each well of the 96 well, with the plate sitting on ice to prevent escape before the plate was full.

The plate was then sealed with gas permeant film. To image the plates over time, they were placed on a circular rotating platform, and moved under a camera to be imaged every hour, with three such platforms or wheels arranged in a stack. At least 200 adults were assayed for genotype, and custom Python scripts used to align the images of each plate and then track the movement of the flies in each well (https://github.com/tjs23/unknome). Lifespan was defined as the time point after the last change in position of the fly in the well. Individual data for both starvation and ROS conditions is in Table S2, with the means and the variances errors plotted in the figures provided in Table S3.

### iFly climbing assay

The climbing speed of flies was measured using the iFly tracking system in which a single camera and mirrors are used to follow the movement of flies in a vial (*75, 76*). The RNAi stocks for the unknome set were crossed to the ubiquitous daughterless-Gal4 driver, and progeny collected at 8 days and 22 days post-eclosion. To follow locomotion, eight flies were placed in a vial which was tapped to collect them at the bottom, and then vial placed in the iFly apparatus for filming over 30 seconds, with this repeated three times. Locomotion velocities were then determined using the iFly tracking software (Kohlhoff et al., 2011). Individual data from both 8 days and 22 days is in Table S2, with the means and the variances errors plotted in the figures provided in Table S3.

### Summary of statistical methods

The general approach we took is as follows, with full details provided below in a Statistical Supplement. We first modelled the distributions of the experimental results relating to each of the phenotypes under consideration parametrically. We thus formalised the goal of identifying outlying genes as identifying outlying sets of parameters corresponding to genes for each of the different phenotypes. Our approach involved three steps. First, we performed a regression to obtain estimates of the parameters for genes and an estimate of their variance– covariance matrix whilst controlling for batch and other effects. This was important because variability across batches was substantial for several of the phenotypes considered. The particular regression model used for this batch correction depended on the dataset.

The next step involved determining an outlier region. To do this, we transformed the parameter estimates so they more closely resembled a sample from a normal distribution such that an elliptical outlier region was appropriate. This transformation was often simply chosen as the identity, but in certain cases logistic transformations were used, for example. To describe how this region was determined, it will be helpful to fix the phenotype and write *μ*_1_, …, *μ*_*J*_for the unknown transformed parameters for the genes, where *J* is the total number of genes under consideration for that phenotype. Furthermore, let us write 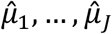 for the corresponding (transformed) estimated parameters. Note that the *μ*_*j*_ were two-dimensional in most examples.

We modelled the *μ*_*j*_ as samples from a mixture of a normal distribution and a distribution of outliers, and aimed to estimate the mean and variance matrix of this normal distribution to give the center and shape of the outlier region. The mean was estimated using a robust mean estimator applied to 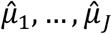 such that the outlying genes did not influence the estimate. Analogously, we also obtained a robust estimate of the variance of the 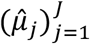 to better reflect the variance of the bulk of the 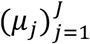. We then employed a bootstrap approach (Efron and Tibshirani, 1994), to adjust this variance estimate to account for the sampling variability of the 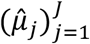 : the raw robust variance would be an overestimate of the corresponding quantity for the true transformed parameters.

Given the final mean and variance estimates, we took our outlier region to be the complement of the elliptical contour of a normal density with this mean and variance with a size such that the probability of falling outside the region was either 0.05 or 0.1, depending on the dataset. Note that in the cases where the parameters *μ*_*j*_ were one-dimensional, the ellipse was simply an interval. Finally, we performed a bootstrap hypothesis test for each gene *j* with the null hypothesis being that *μ*_*j*_ falls within the outlier ellipse. We thus obtained *p*-values for each gene quantifying the evidence that it is an outlier according to the data. Note that this measure incorporates both how outlying 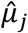 is, but importantly also takes into account the fact that 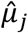 is a noisy estimate of the true *μ*_*j*_. These *p*-values were then corrected for multiple testing using the Benjamini–Hochberg procedure (Benjamini and Hochberg, 1995).

### CRISPR/Cas9-mediated knock-out

CRISPR target sites were chosen using the CRISPR Optimal Target Finder (http://targetfinder.flycrispr.neuro.brown.edu/). pCFD3 was used for BbsI-dependent gRNA cloning (http://www.crisprflydesign.org/) (Port et al., 2014). gRNA transgenics were generated for all candidate genes using BDSC stocks #25709 or #25710, depending on the chromosomal location of the target gene. To generate indels transgenic gRNA lines were crossed to either CFD2 or TH_attP2. DNA microinjections were performed by the University of Cambridge Department of Genetics Fly Facility. For generation of *CG10795* mutants, gRNAs were cloned into pCFD3, and plasmids injected into CFD2 embryos. Stable stocks were generated to recover indels for all candidate genes. For genotyping, single males were collected and the genomic DNA was isolated using microLYSIS-Plus (Clent Life Science). Diagnostic PCRs followed by sequencing identified indels.

### F**ertility assays on CRISPR/Cas9 mutants**

To check male fertility, crosses with 5 Oregon R wild-type virgins and 3 mutant males were set up for each genotype. Crosses were kept at 25°C and knocked over twice. The total number of offspring was counted for all crosses and the mean +/-SD was plotted for each genotype. Deficiencies uncovering the candidate genes were used to check for potential off-target effects. To check female fertility three crosses with 5 mutant virgins and 3 Oregon R wild-type males were set up for each genotype, and processed in the same was as for male fertility. A deficiency uncovering *CG8237* was used to check for potential off-target effects.

### Analysis of *CG11103* and *CG10795* embryonic phenotypes

Overnight egg collections (at 25°C) from *CG11103* and *CG10795* mutant females and males were kept at 25°C for 48 hours. Dead embryos were dechorionated and mounted in Hoyer’s Medium. Slides were kept at 65°C for at least 24 hours and widefield amges obtained with a Zeiss Axioplan. For examination of Elav expression, overnight egg collections from *CG11103* and *CG10795* mutant females and males were dechorionated with bleach and fixed using 4% formaldehyde. Embryos were devitellinised using n-Heptane/Methanol. Embryos were washed in PBT 0.1% Tween20 and blocked in PBT 0.1% Tween20 plus 5% BSA. Mouse anti-Elav (1/20; DSHB), were added over night at 4°C, and then embryos washed in PBT 0.1% Tween20. Donkey anti-mouse Alexa 488 (Fisher Scientific) was added and left for 2h at RT. Embryos washed in PBT 0.1% Tween20, and mounted in Vectashield containing DAPI (Vector Laboratories) and imaged on a Zeiss LSM 710 confocal.

### Analysis of male seminal vesicles in *CG6153* mutants

Testes from 3-5 days old adult males were dissected in PBS and then either directly transferred onto a slide with Schneider’s medium to take live images using a Zeiss 710 confocal microscope or fixed in 4% paraformaldehyde for 30 min at RT. PFA was then removed and the testes washed in PBT 0.1% Tween 20. Images were taken on a Zeiss stereo-microscope and a Nikon digital camera.

### Statistical Supplement

Here we provide further details to complement the outline of the statistical methods presented in the STAR Methods section. The approach used consists of three steps: modelling the data to associate each gene with statistical parameters; construction of an outlier region in the space of these parameters; and performing hypothesis tests to determine whether these parameters fall within this region. The material here is similarly organised into three sections describing each of these steps. In much of what follows, we consider the experimental setting (i.e. fertility, wing size etc.) to be fixed and describe general procedures that we apply, with some modifications, to each of the settings.

## 1 Modelling the data

### 1.1 General procedure

The data for each experiment takes the general form (*Y*_*ij*_, *x*_*ij*_) ∈ℝ^*d*^ × ℝ^*p*^, *i* = 1, …, *n*_*j*_, *j* = 1, …, *J*, where *J* was the total number of genes, and *d* ∈{1,2}. Here *Y*_*ij*_ corresponds to the *i*th measurement taken on the *j*th gene and the *x*_*ij*_ are associated covariates that may indicate the batch in which the measurement was taken, for example. Our goal is to identify outlying genes, and for this purpose we first construct a parametric model for the data of the form

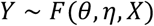

where *Y* and *X* collect together the response of covariates respectively, *θ* = (*θ*_1_, …, *θ*_*J*_) ∈ℝ^*d*×*J*^ are the parameters associated the with genes and *η* represents a collection of nuisance parameters (e.g. parameters associated with the different batches). The statistical problem at hand then is to identify outlying *θ*_*j*_. For this we need to introduce a notion of what it means to be an outlier, and then propose a methodology for testing for each *j* whether *θ*_*j*_ is an outlier. These latter two tasks are described in Sections 2 and 3. To do these, we require estimates 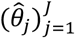 of 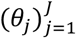 that are approximately unbiased and Gaussian with estimated variance 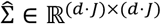. Note that as we are only interested in differences between different *θ*_*j*_, we are for example free to introduce a sum-to-zero constraint on these parameters to reduce the overall variance, and we do this throughout.

Below we present the specific statistical models *F* used for each of the different experimental datasets for which (versions of) maximum likelihood estimation then delivers these quantities. All computations were performed in R (R Core Team, 2018).

### 1.2 Fertility

Let *Y*_*ijk*_ ∈ℤ be the *i*th brood size measurement corresponding to female flies with gene type *j* in batch *k* for *i* = 1, …, *n*_*ji*_ (where *n*_*jk*_ may be 0 for some (*j, k*)). We will first present our analysis of the data on females; analysis for the data on males proceeded similarly.

To examine the mean–variance relationship (i.e. how 𝔼(*Y*_*ijk*_) relates to Var(*Y*_*ijk*_)), we first formed for all (*j, k*) such that *n*_*jk*_ ≥ 2,

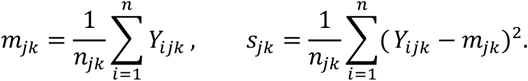

We then regressed *m*_*jk*_ on to *s*_*jk*_ via the following optimisation:

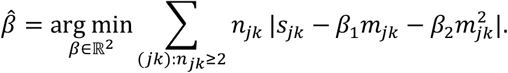

The lack of intercept in this regression encodes the restriction that when 𝔼(*Y*_*ijk*_) = 0 we must have Var(*Y*_*ijk*_) = 0; the use of the absolute value rather than the more usual squared error loss is to account for the exponential-type tails we may expect for the *s*_*jk*_; and the weights *n*_*jk*_ reflect the variance of the *s*_*jk*_.

We thus obtained an estimated variance function 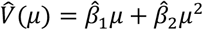 such that 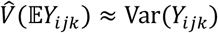 We obtained coefficients

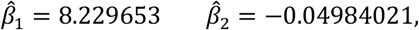

and as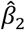 was negative, we were able to express 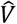 as a scaled version of a Bernoulli variance function via

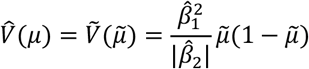

with 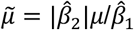 To fit a regression model with this form of variance function, we used a quasi-binomial regression after transforming the data 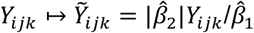 The transformed data took values in [0,1] so we used a logit link and modelled the mean 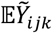 as

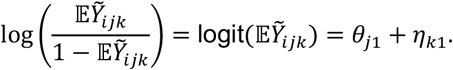

To handle zero counts we used the bias correction of (Firth, 1993), as implemented in (Kosmidis, 2019), which always produces finite parameter estimates. The analysis of the male data was very similar, and in the end we obtained estimates 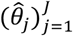 and block diagonal estimated variance matrix 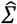 (as the male and female data were independent).

### 1.3 Wing size

Let *Y*_*ijk*_ ∈ℝ^2^ be the *i*th measurement on the *j*th gene in the *k*th batch, defined for *i* = 1, …, *n*_*jk*_ (where *n*_*jk*_ may be 0 for some (*j, k*)) with first and second components denoting measurements for anterior and posterior wing segments respectively. We used the model

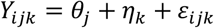

where 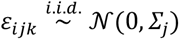 with Σ ∈ℝ^2×2^. Inspection of the data showed that the correlation matrices corresponding to the Σ_*j*_ vary very little over *j* and the difference is barely detectable by permutation tests. We therefore constrained Σ_*j*_ in the following way: 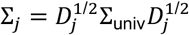 where Σ_univ_ ∈ℝ^2×2^ is a universal correlation matrix and the *D*_*j*_ ∈ℝ^2×2^ are diagonal matrices with variances corresponding to each gene.

### 1.4 PolyQ aggregates

We first constructed from the available data two quantities from each replicate: the number of aggregates with area in pixels greater than 50, and the corresponding number with area less than or equal to 50. They form the components of *Y*_*ijk*_ ∈ℤ^2^, for which we use quasi-Poisson models with log links as follows.

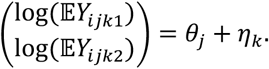

We performed two separate Poisson regressions for each component of the response. In order to avoid issues where parameter estimates from standard maximum likelihood estimation were too large, we employed the bias correction of (Firth, 1993), as implemented in (Kosmidis, 2019). To estimate the covariance matrix of the parameters, we noted that the working residuals from the regressions displayed a covariance that was constant across fitted values from each of the regressions. Using this estimated covariance and estimated dispersion parameters we formed a full covariance matrix 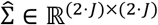 for all 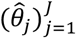

### 1.5 Survival under stress

Let *Y*_*ijkl*_ and 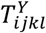 denote the censored survival and censoring times under oxidative stress for the *i*th replicate of gene *j* in batch *k* and wheel *l*. We fitted a Cox proportional hazards model of the form

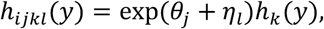

where *h*_*ijkl*_ is the hazard function of the unobserved uncensored version 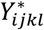 of *Y*_*ijkl*_, and *h*_*k*_ is an unspecified baseline hazard function for batch *k*.

We used an analogous model for the data concerning survival times under starvation.

### 1.6 Climbing speed

Let *Y*_*ijkl*_ and *Z*_*ijkl*_ denote the *i*th speed measurement corresponding to gene *j*, batch *k* and repeat *l* for days 8and 22 respectively. We used the following random effects models:

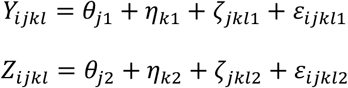

where 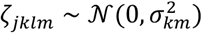 and 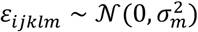 all independently.

## 2 Outlier region construction

From the initial regression, we obtained estimates 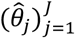 for the parameters 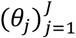 corresponding to each gene, and their associated estimated variance matrix 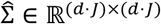. In order that an elliptical outlier region was appropriate, we transformed the estimates depending on their distribution to 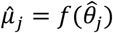 where the transform function *f*: ℝ^*d*^ → ℝ^*d*^ chosen is given in Table 1.

**Table 1.** Transform functions used for different datasets.

| Dataset | Transform function $f$ |
| --- | --- |
| Fertility | logistic |
| Wing size | identity |
| PolyQ aggregates | exponential |
| Survival under Stress | identity |
| Climbing speed | identity |

Let us write *μ*_*j*_ = *f*(*θ*_*j*_) for each *j* = 1, …, *J*. We considered *f* as fixed, and as is common in the analysis of outliers, considered a model for the *μ*_*j*_ as samples from a mixture of a normal distribution and an outlier distribution *F*_out_ (Hawkins, 1980):

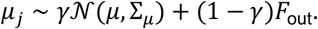

We assumed the mixture proportion *γ*to be greater than 0.5 and that the support of *F*_out_ was sufficiently far from *μ*. We used the minimum covariance determinant estimator (Rousseuw and Driessen, 1999), as implemented in (Maechler et al., 2018), to give a robust estimate 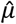 of *μ* and an initial estimate 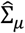 of Σ_*μ*_. Whilst we can expect that 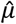 is a reasonable estimate of 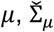 will be substantially inflated by the sampling variability of the 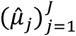. To correct for this, we employed the following bootstrap strategy.

1. Produce bootstrap samples 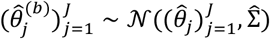 for *b* = 1, …, B.
2. Form 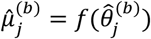 for *j* = 1, …, *J* and *b* = 1, …, B.
3. Compute robust covariance estimates 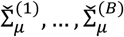 based on each of the bootstrap samples 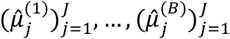 using the minimum covariance determinant estimator.
4. Set

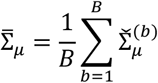

and finally define our final estimate 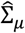 of Σ_*μ*_ by

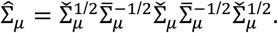

The rationale for this approach is that

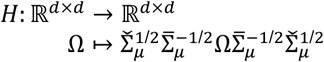

is a mapping that satisfies 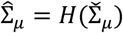 and

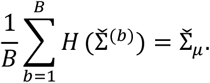

Thus, we can think of *H* as a corrective transformation that were 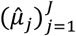 to be a sample from the ground truth, gives an approximately unbiased estimate of its (robust) covariance. Applying *H* to 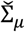 should similarly correct it to give a better estimate of Σ_*μ*_. The reason for generating the bootstrap samples at the level of the untransformed parameters is that the Gaussian approximation in step 1 of the procedure above, which mimics the sampling distribution of the 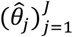, would typically be more reliable than the analogous approximation for the 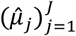.

Given our final estimates 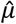 and 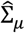 we set the outlier region to be the complement of the elliptical contour of a 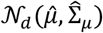 density such that the probability of 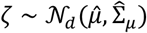 falling within the region is given by 0.05 or 0.1, depending on the dataset. This outlier region can be mapped to the *θ*-space using the inverse of *f*; in the sequel we will refer to this region as *R*.

## 3 Testing for outliers

Given outlier region *R* such that all *j* for which *θ*_*j*_ ∈*R* are deemed outliers, we constructed for each *j* an (approximate) *p*-value *p*_*j*_for the null hypothesis *θ*_*j*_∉*R*. In the cases where the region was an interval, this was straightforward. In the cases where the region was two-dimensional, this was done using a bootstrap scheme, the main steps of which were as follows. Denote by *A* the complement of *R*, and also let *A*_*μ*_ be the (elliptical) region *f*(*A*).

1. Compute via the delta method an estimate 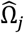 of the variance of 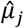.
2. Compute the projection 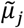 of 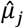 on to the elliptical region *A*_*μ*_ using the Mahanolobis distance with covariance 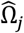 :

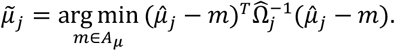 (Details for how this is performed are given in Section 4.)
3. Set

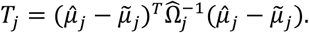 Also define 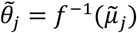.
4. Let 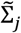 be an estimate of the maximum likelihood estimate of *θ*_*j*_under the null that *θ*_*j*_∈*A*. Generate *B* = 100000bootstrap samples 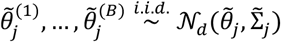. Let 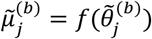
5. Compute bootstrap versions of the test statistic *T*_*j*_:

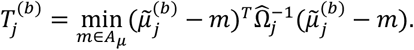
6. Then

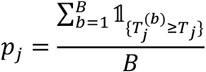

in a Monte Carlo estimate of the *p*-value. To improve the quality of this estimate, we in fact used an importance sampling scheme where initially the 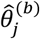 were generated from a mixture of the Gaussian distribution above, and 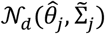 (with mixture proportions 0.5); the 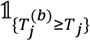 terms were then weighted according to the importance sampling weights.

The rationale for this is as follows. The test statistic *T*_*j*_ encapsulates how far 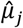 is from the region *A*_*μ*_ taking into account the variance of the 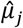 (directions in which 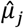 is highly variable are effectively down-weighted). Under the null hypothesis that *θ*_*j*_ ∈*A*, we should have 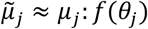 and so the bootstrap distribution should approximate the null distribution and thus provide effective calibration for *T*_*j*_.

We finally apply false discovery rate (FDR) correction to the *p*-values using the Benjamini– Hochberg procedure (Benjamini and Hochberg, 1995). Although controlling for batches and the fact that the outlier region is determined using the data would make the *p*-values dependent, the dependence should be weak and thus the Benjamini–Hochberg procedure should at least approximately control the FDR.

## 4 Ellipse projection

Here we describe an efficient approach to computing

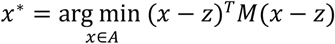

where *M*∈ℝ^*d*×*d*^ is a symmetric positive definite matrix and ellipsoid *A* = {*x*: *x*^*T*^Ω*x* ≤ *c*} for symmetric positive definite Ω ∈ℝ^*d*×*d*^, *c* > 0 and *z* ∉*A*. Equivalently, the problem is to find the minimum *c*^∗^ > 0 such that there exists *x*^∗^ with

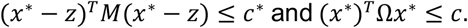

By Lagrangian duality, we know there exists *λ* that

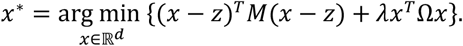

Consider the eigendecomposition *M* = *PD*^2^*P*^*T*^. Writing *y*^∗^ = *DP*^*T*^*x*^∗^ we have

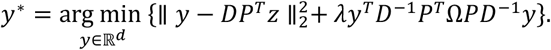

Let the eigendecomposition of *D*^−1^*P*^*T*^Ω*PD*^−1^ be *U*Λ*U*^*T*^. We see that then

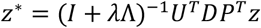

where *z*^∗^ = *U*^*T*^*y*^∗^ so *x*^∗^ = *PD*^−1^*Uz*^∗^ and *λ* is such that (*z*^∗^)^*T*^Λ*z*^∗^ = *c*.

